# Enhanced auxin signaling hub triggers root hair growth at moderate low temperature in *Arabidopsis thaliana*

**DOI:** 10.1101/2024.08.15.608078

**Authors:** Victoria Berdion Gabarain, Gerardo Núñez-Lillo, Aleš Pěnčík, Miguel Angel Ibeas, Javier Martínez Pacheco, Leonel Lopez, Mariana Carignani Sardoy, Jia Zhongtao, Ondřej Novák, Ricardo F.H. Giehl, Nicolaus von Wiren, Claudio Meneses, José M. Estevez

**Affiliations:** Fundación Instituto Leloir and IIBBA-CONICET. Av. Patricias Argentinas 435, Buenos Aires C1405BWE, Argentina; Escuela de Agronomía, Facultad de Ciencias Agronómicas y de los Alimentos, Pontificia Universidad Católica de Valparaíso, Calle San Francisco s/n, La Palma, Quillota 2260000, Chile; Laboratory of Growth Regulators, Faculty of Science of Palacký University & Institute of Experimental Botany of the Czech Academy of Sciences, Šlechtitelů 27, CZ-779 00 Olomouc, Czech Republic; ANID - Millennium Science Initiative Program - Millennium Nucleus for the Development of Super Adaptable Plants (MN-SAP), Santiago, Chile; ANID - Millennium Science Initiative Program - Millennium Institute for Integrative Biology (iBio) Santiago, Chile; Centro de Biotecnología Vegetal, Facultad de Ciencias de la Vida, Universidad Andres Bello, Santiago, Chile; College of Resources and Environmental Sciences, National 13 Academy of Agriculture Green Development, China Agricultural University, 14 100193 Beijing, China; Molecular Plant Nutrition, Department of Physiology & Cell Biology, Leibniz Institute of Plant Genetics and Crop Plant Research (IPK), Corrensstr. 3, 06466 Gatersleben, Germany; Departamento de Fruticultura y Enología, Facultad de Agronomía y Sistema Naturales, Pontificia Universidad Católica de Chile, Santiago, Chile; Departamento de Genética Molecular y Microbiología, Facultad de Ciencias Biológicas, Pontificia Universidad Católica de Chile, Santiago, Chile; ANID - Millennium Science Initiative Program - Millennium Institute, Center for Genome Regulation, Santiago, Chile

**Keywords:** Arabidopsis, auxin, low temperature, root hairs

## Abstract

Root hairs (RH) as a mixed tip- and non-tip growing protrusions that develop from root epidermal cells are important for nutrient and water uptake, root anchoring, and interaction with soil microorganisms. Although nutrient availability and temperature are critical interlinked factors for sustained plant growth, the molecular mechanisms underlying their sensing and downstream signaling pathways remain unclear. Here, we identified a moderate low temperature (10°C) condition that triggers a strong RH elongation response involving several molecular components of the auxin pathway. Then, we have determined that auxin biosynthesis carried out by YUCCAs/TAA1, the auxin transport conducted by PIN2/PIN4 and AUX1/PGP4, and the auxin signaling controlled by TIR1/AFB2 coupled to four specific ARFs (ARF6/ARF8 and ARF7/ARF19), are all crucial for the RH response at moderate low temperature. These results uncover the auxin pathway as one central hub under moderate low temperature in the roots to trigger RH growth. Our work highlights the importance of moderate low temperature stimulus as a complex nutritional signal from the media soil into the roots that may be fine-tuned for future biotechnological applications to enhance nutrient uptake.

## Introduction

Root hairs (RH) as mixed tip and non-tip-growing protrusions that develop from root epidermal cells are important for nutrient, water uptake as well as for root anchoring and interaction with soil microorganisms (Balcerowick et al. 2015; Herburger et al. 2023). RH development involves cell fate determination, RH initiation, and then cell elongation by polarized or tip growth. In Arabidopsis (*Arabidopsis thaliana*), RH cells (T, trichoblasts) and non-hair cells (AT, atrichoblast) files differentiate from the epidermal cells in the meristematic and transition zones of the root (Dolan *et al*., 1993). RH constantly integrates inner signals such as hormones (e.g. auxin and ethylene) and environmental cues like nutrients (e.g., phosphates and nitrates) and temperature (e.g., low temperature) (Balcerowick *et al*. 2015; Shibata *et al*. 2019; Kia *et al*. 2022: Lopez et al. 2023). The auxin effect in RH growth depends on systemic auxin metabolism focused mainly on the root meristem-root cup, intercellular auxin transport via root epidermis (Sauer *et al*., 2013) and on site RH signaling (Casal & Estevez, 2021; Lopez *et al.,* 2023). In recent years, it has become evident that RH play a crucial role in root activities, making them a potential target for biotechnological enhancement in commercial cultivars. The integration of hormonal pathways including auxin with environmental signals and the fine-tuning of RH development in plants are intricate and poorly understood processes.

The main natural auxin, indole-3-acetic acid (IAA), is biosynthesized primarily via two chemical reactions (Zhao, 2012). The amino acid L-tryptophan (Trp) is produced through the shikimate pathway and is the primary precursor in plants’ four major auxin biosynthesis pathways. These pathways are known as the indole-3-acetamide (IAM), indole-3-acetaldoxime (IAOx), tryptamine (TRA), and indole-3-pyruvic acid (IPyA) pathways, named after their respective major intermediate compounds (Mano and Nemoto, 2012; Ljung, 2013; Kasahara, 2016). The major sources of newly produced auxin are thought to be these routes. However, there may also be Trp-independent mechanisms (Tivendale *et al*., 2014; Wang *et al*., 2015). After being synthesized and transported, auxin may undergo degradation through conjugation and subsequent oxidation (Hayashi et al, 2021). This process results in the formation of two main metabolites: 2-oxindole-3-acetic acid (oxIAA) and oxIAA-glucose (oxIAA-Glc) (Östin *et al*., 1998; Kai *et al*., 2007; Pěnčík *et al*., 2013). Inactive auxin involves conjugates with amino acids or sugars (Tam *et al*., 2000; Kowalczyk and Sandberg, 2001). Specific conjugates may undergo hydrolysis, resulting in the release of active auxin. This suggests that these conjugates might function as temporary storage forms of the inactive hormone (Ludwig-Müller 2011). However, in *Arabidopsis thaliana*, the most prevalent amide-linked conjugates, IAA-aspartate (IAA-Asp) and IAA-glutamate (IAA-Glu), do not undergo or it is very low the reversible conversion to IAA. Instead, they likely function as degradation intermediates (Kowalczyk and Sandberg, 2001; Woodward and Bartel, 2005). IPyA is converted to IAA by the catalysis of flavin-containing monooxygenases encoded in YUCCA (YUC) genes (Zhao *et al*., 2001; Mashiguchi *et al*., 2011; Won *et al*., 2011). Downstream biosynthesis, various auxin transporters regulate cellular auxin levels including the auxin efflux carriers PIN-FORMED (PIN) and ATP-BINDING CASSETTE B (ABCB) as well as the auxin influx carriers from the AUXIN RESISTANT1 (AUX1)/LIKE AUX1 (LAX) family. PIN2 is the only family member expressed in root epidermal cells and is present in both root and non-RH cells (Luschnig *et al*., 1998), with higher PIN2 abundance in atrichoblasts (Löfke *et al*., 2015). PIN2 displays a polar localization to the apical (shoot-ward) side of the epidermal cells and, thereby, mediates the basipetal (shoot-ward) direction of the auxin flow (Wisniewska *et al*., 2006; Cho *et al*., 2007; Rigas *et al*., 2013). Members from the AUX1/LAX protein family localize to the plasma membrane and mediate auxin uptake into the cell (Yang *et al*., 2006; Swarup & Péret, 2012). The H^+^/IAA symporter AUX1 was demonstrated to facilitate auxin influx (Yang *et al*., 2006; Yang & Murphy, 2009; Ikeda et al. 2009). AUX1 overexpression enhances RH growth, presumably due to the increased auxin levels in the root hair cells (Ganguly *et al*., 2010). Intriguingly, in the root epidermis of wild-type plants, AUX1 is expressed only in non-hair cells (Jones *et al*., 2009), suggesting that auxin transport in non-hair cells sustains RH development and cell elongation (Lee & Cho, 2006; Jones *et al*., 2009; Ikeda et al. 2009). Finally, the slow genomic auxin response is mediated by the receptors of the TRANSPORT INHIBITOR RESPONSE1/AUXIN SIGNALING F-BOX (TIR1/AFB) family (Dharmasiri *et al*., 2005; Kepinsky & Leyser, 2005). Auxin binding on TIR1/AFB and its coreceptor AUXIN RESISTANT/INDOLE-3-ACETIC ACID INDUCIBLE (AUX/IAA) induces the proteasome-dependent degradation of the latter. AUX/IAA depletion results in the subsequent release of ARF TFs, which modify the auxin-dependent expression profile. Once in the trichoblasts, auxin sensed *in situ* by the TIR1/AFBs and Aux/IAA coreceptors promotes RH growth. ARFs control the expansion of RH (Mangano *et al*., 2017; Bhosale *et al*., 2018; Schoenaers *et al.,* 2018; Jia *et al.,* 2023). ARF7 and ARF19 are the most abundantly expressed ARFs in root hair cells (Bargmann *et al*., 2013) and they were shown to be linked to low Pi conditions (Bhosale *et al.,* 2018; Giri et al., 2018; Schoenaers *et al*., 2018). On the other hand, ARF6 and ARF8 were shown to be involved in the response of RHs to low nitrate (Jia *et al.,* 2023). Downstream of ARFs, auxin treatments promote RH growth by positively regulating the expression of several TFs, including the main RH regulators bHLH RHD6 and RSL4 (Yi *et al*., 2010) and several other TFs such as RSL2 and LRL3 (Karas *et al*., 2009; Pires *et al*., 2013).

When exposing Arabidopsis plant accessions to moderate low temperature condition (around 10°C), an enhanced RH growth (up to three-fold) was detected (Moisön *et al*., 2021; Pacheco *et al*., 2022; Pacheco *et al.,* 2023) although the rest of the plant and roots arrested their growth. This unexpected and specific growth response in RH raised several questions regarding the molecular mechanisms behind it. It was hypothesized that a low availability and mobility of nutrients caused by moderate low temperature might be the direct stimulus that promotes RH growth (Moisön *et al*. 2021; Pacheco et al. 2021). Recently, a link between the nitrate availability and this specific low temperature growth was found (Pacheco et al. 2023), and, a low nitrate condition promotes an active RH growth mediated by some auxin pathway components (Jia et al. 2023). In addition, it was shown that low phosphate also triggers an upregulation of auxin biosynthesis in the tip of the root with a concomitant enhancement of RH growth by using other molecular components than low nitrate (Bhosale et al. 2018; Jia et al. 2023; Lopez et al. 2023). It is known that downstream auxin, ROOT HAIR DEFECTIVE 6 (RHD6) and RHD6-like 4 (RSL4) are the main transcription factors that regulate RH growth in low phosphate (Bhosale et al.2018), low nitrate (Jia et al. 2023), and low temperature conditions (Moison et al. 2021). Previously, it was shown that the long non-coding RNA APOLO identifies the locus encoding for the master regulator (RHD6) and regulates RHD6 transcriptional activity, which in turn causes moderate low temperature-induced RH elongation by activating the RSL4 transcription factor gene (Moison et al. 2021). Additionally, it was shown that APOLO modifies transcription factor WRKY42’s binding to the RHD6 promoter through interactions with it. Moderate low temperature activates RHD6 only when WRKY42 is present, and WRKY42 deregulation prevents cold-induced RH expansion. All of our findings point to the formation of a new ribonucleoprotein complex including APOLO and WRKY42 that functions as a regulatory hub to shape the epigenetic environment of RHD6 and integrate signals controlling RH growth and development (Moison et al. 2021). In this work, we question if the moderate low temperature condition can trigger changes in auxin synthesis, transport, and signaling, as well as the molecular components involved in the roots as a consequence of the effect on the mobility of the nutrients in the media-soil. Using several diverse approaches, we identified the precise role of the auxin pathway elements during RH polar growth in moderate low temperature conditions. We found that mutant and overexpression lines affected auxin biosynthesis, transport, and downstream signaling, altering the ability to respond to moderate low temperature at the RH level. Our study has not only identified the detailed molecular components of the auxin pathway that govern how RH grows under moderate low temperature, but also highlights possible new targets to be reprogrammed in crop engineering to enhance nutrient uptake by the roots.

## Results

### Low temperature induces significant transcriptional changes in the auxin pathway in roots

Root hair (RH) development is a well-conserved adaptation to low soil nutrients. Previous research indicates that growth at 10°C decreases nutrient availability in the growth medium, which has a major impact on RH growth. Pacheco *et al*. (2022) found that this effect limits nitrate transport and accessibility in culture medium, promoting polar RH growth. Previously, auxin was linked to both low nitrate (Jia *et al.,* 2023) and low phosphate promotion of RH growth (Giri *et al*., 2018; Bhosale *et al*., 2018). This evidence allowed us to use the moderate low temperature effect on RH to uncover the molecular mechanisms linked to complex nutritional cues. We then asked if the auxin pathway could be involved in moderate low temperature mediated RH growth. To explore auxin-related genes, we first identified which were the Arabidopsis genes annotated in the different areas of the auxin pathway: biosynthesis, conjugation, signaling and transport. A total of 349 auxin-related genes based on annotation (**Table S1**) were then used to evaluate their expression at 10°C for 2h and 6h in Wt Col-0 as well as in the hairless double knockout mutant *rsl2 rsl4* (**Figure 1**). RSL2 and RSL4 are major TFs that control cell elongation in RH at moderate low temperature (Moisön *et al*., 2021). Differential expression analysis compared 2h and 6h of low temperature treatment to basal condition at 22°C in Col-0 genotype to identify low temperature-response genes. A total of 7,958 and 12,860 differentially expressed genes were identified at 2h and 6h of cold treatment, respectively. Using a list of 318 annotated auxin-related genes (**Table S1**), we identified a total of 170 differentially expressed auxin-related genes in the moderate low temperature treatments (**Table S2**). The expression values of the 170 auxin-related genes with differential expression were scaled and clustered in the heat-map of **Figure 1A**. Four clusters were identified, cluster 1 (49 DEG), which contains PIL2,5,6 and TIR1 and cluster 2 (29 DEG), which contains genes downregulated by low temperature treatment. Cluster 3 (22 DEG) contains TAA1, ARF19, PIN1,PIN4 and cluster 4 (70 DEG) contains YUCCA genes (YUCCA4,5,6,8), several ARFs (ARF6,7,8), several IAAs (IAA14,17,18,28), few PILs (Barbez et al. 2012) and PIN2 that are upregulated by low temperature treatment. On the other hand, in the *rsl2 rsl4* double mutant the genes of cluster 2 and cluster 4 have significantly higher expression values than Col-0 genotype suggesting a negative regulation. In comparisson, genes of cluster 3 have substantially lower expression values than the Col-0 genotype, possibly by a positive regulation by RSL2-RSL4 (**Figure 1B**). A comparative gene ontology analysis was conducted to understand which biological processes are enriched in the 4 clusters identified before (**Figure 1C**). Cluster 1 presents GO terms associated with auxin transport and regulation of growth. In constrast, in cluster 2, GO terms related to root morphogenesis and amine biosynthetic process and auxin metabolic process can be found. In cluster 3 GO terms of gravitropism, auxin polar transport and auxin biosynthetic process are highlighted. Finally, in cluster 4, GO terms associated with root morphogenesis, meristem growth and regulation of the flavonoid biosynthetic process are observed. This data shows that auxin biosynthesis, transport, and signaling-related genes are globally affected by the moderate low temperature treatment in roots.

**Figure 1.**
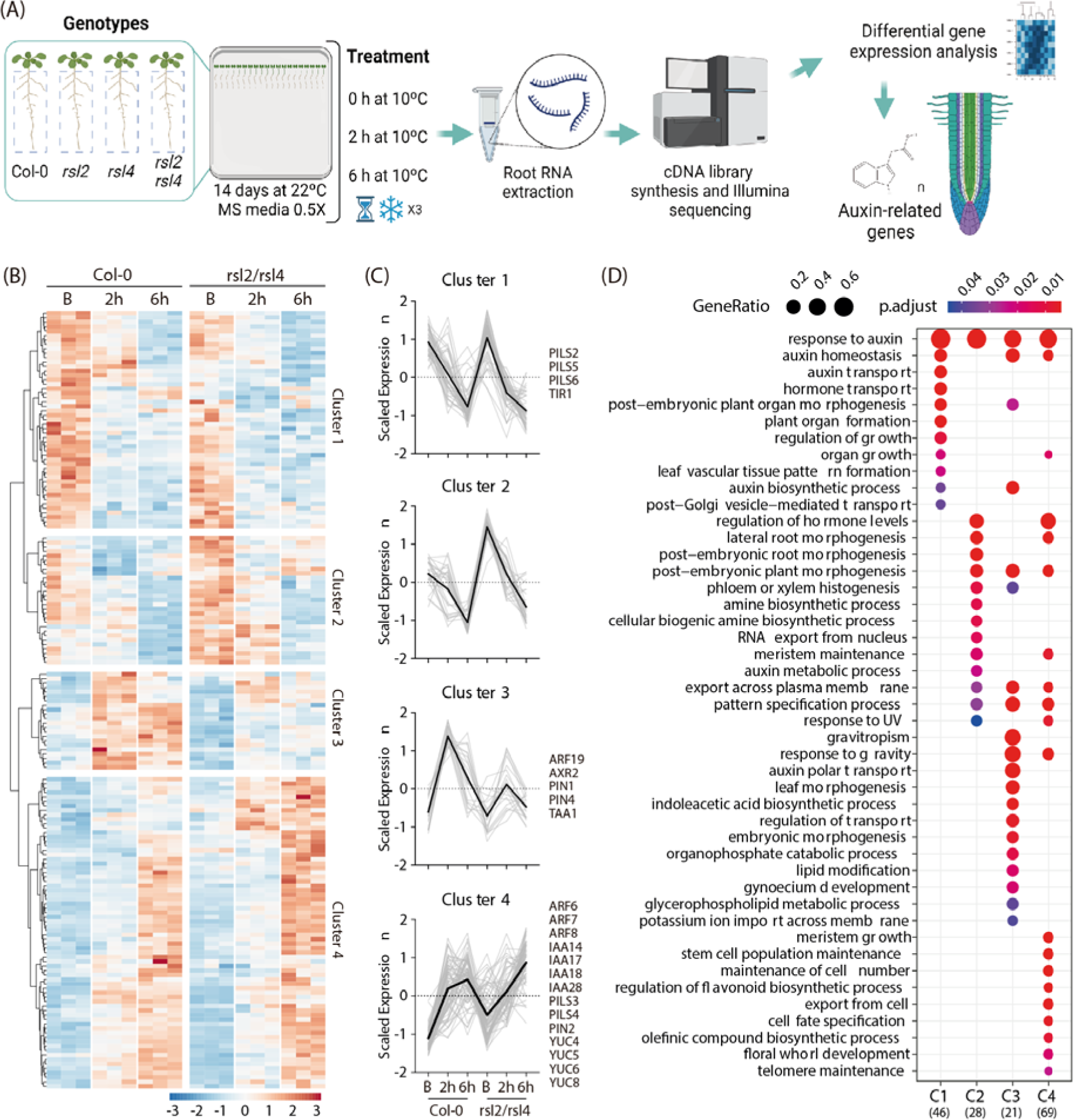
Moderately low temperatures trigger global transcriptomic changes in the auxin pathway. (**A**) An scheme with the RNA-seq procedure was followed to identify auxin-related genes at moderate low temperature. **(B)** Heatmap showing the hierarchical gene clustering for 170 auxin-related genes differentially expressed between room temperature (22°C) and low temperature (10 °C) growth roots in wild type (Col-0) and *rsl2 rsl4* double mutant. Gene expression values were scaled considering the mean centered, divided by standard deviation, and represented in a blue-red color scale. Clustering analysis was made using the ward.D2 method. **(C)** Scaled expression values of each auxin-related gene (gray lines) and the tendency of each cluster (black line) were graphed. Most representative genes were listed on the right side of each corresponding cluster. **(D)** Gene ontology term enrichment analysis of genes belonging to each one of the identified 4 Clusters (C1-C4). Vertical lines represent the information of each identified cluster while the horizontal lines represent the enriched GO terms. The numbers at the bottom correspond to the number of genes used in each cluster. The blue-red scale color represents the adjusted p-value, and the point size represents the gene ratio. See also **Table S1**.

### Auxin biosynthesis in the root tip is required for RH growth at a moderate low temperature

Auxin is essential for RH growth and development under variable conditions (Mangano *et al*., 2017; Bhosale *et al*., 2018; Jia *et al*., 2023). Tryptophan aminotransferases of the TRYPTOPHAN AMINOTRANSFERASE OF ARABIDOPSIS (TAA)/TRYPTOPHAN AMINOTRANSFERASE RELATED (TAR) family and YUCCA-type flavin monooxygenases catalyze the conversion of tryptophan to auxin, representing the major pathway for local auxin biosynthesis in Arabidopsis (Zhao *et al*., 2001). Interestingly, our RNA-seq gene expression profiling revealed that transcript levels of TAA1 and several YUCCA genes increase rapidly in response to low temperature (**Figure 1**). Th RH growth of loss-of-function *taa1* and quintuple *yuc* mutants (*yucq*, lacking *yuc3, yuc5, yuc7, yuc8* and *yuc9*) (Chen *et al*., 2014; Gaillochet *et al*., 2020) were analyzed to functionally validate the role of local auxin biosynthesis in stimulating RH elongation under low temperature. RH lengths of *taa1* and *yucq* mutants were much shorter to the WT at both 22°C and at moderate low temperature (**Figure 2A**). Adding 100 nM of auxin (IAA, indole acetic acid) exogenously, restored both *taa1* and *yucq* mutants to WT Col-0 RH growth (**Figure 2A**). In addition, we select one of the YUCCA genes that are induced by moderate low temperature in the RNA-seq data (see Cluster 4, **Figure 1**) to study in more detail. The overexpression of YUC8 (YUC8-OE) showed much longer RH than WT Col-0 at both temperatures (**Figure 2B**) and YUC8-GFP translational reporter, that catalyzes the downstream reaction of TAA1 in auxin synthesis, showed significantly higher signal under low temperature (**Figure 2C**). To investigate whether auxin biosynthesis mediated by TAA1-YUCCs is involved in the RH elongation response at low temperature, we treated Wt Col-0 plants with the auxin biosynthesis inhibitor yucasin (Nishimura *et al*., 2013) (**Figure 2D**). When yucasin was applied to roots remarkably decreased RH elongation at 22°C (IC50=6 uM) while higher concentrations (IC50=33 uM) were required to block RH growth at low temperature. These results suggest that RH elongation at low temperature relies on local auxin biosynthesis in roots. TAA1-GFP translational reporter showed that protein levels decreased in root apices grown under low temperature after 3 days of treatment (**Figure 2E**). All these results together confirm the role of TAA1 and multiple YUC-dependent auxin synthesis is required to regulate RH elongation under low temperature (**Figure 2F**). Then, we tested if auxin precursor and related metabolites levels were changed under low temperature in the roots of Wt Col-0 and YUC8 OE line. Seedling roots samples were isolated, and auxin precursors and metabolites were identified and quantified using high performance liquid chromatography coupled with mass spectrometry (UHPLC-MS/MS). Free IAA, auxin precursors, related metabolites and auxin conjugates were quantified (**Figure 3** and **Table S3**). After moderate, low temperature treatment, the amount of Trp (Tryptophan) in the roots is 3-fold increased in Wt Col-0. Similar trends are observed for the alternative route IAM and IPyA, that both produce IAA, are drastically enhanced under moderate low temperature in Wt Col-0. In addition, the content of oxIAA, oxIAA-Glu and oxIAA-Glc were also increased by low temperature treatment in Wt Col-0 and specially in the YUC8 OE line (**Figure 3**). This suggests that the excess of IAA might be inactivated in the YUC8 OE. Overall, these results indicate that major changes occur in the auxin pathway when roots are exposed to moderate low temperature. This agrees with the overall transcriptional changes in the auxin-related genes detected (**Figure 1**) and specifically in the biosynthetic genes TAA1-YUCs (**Figure 1** and **2**).

**Figure 2.**
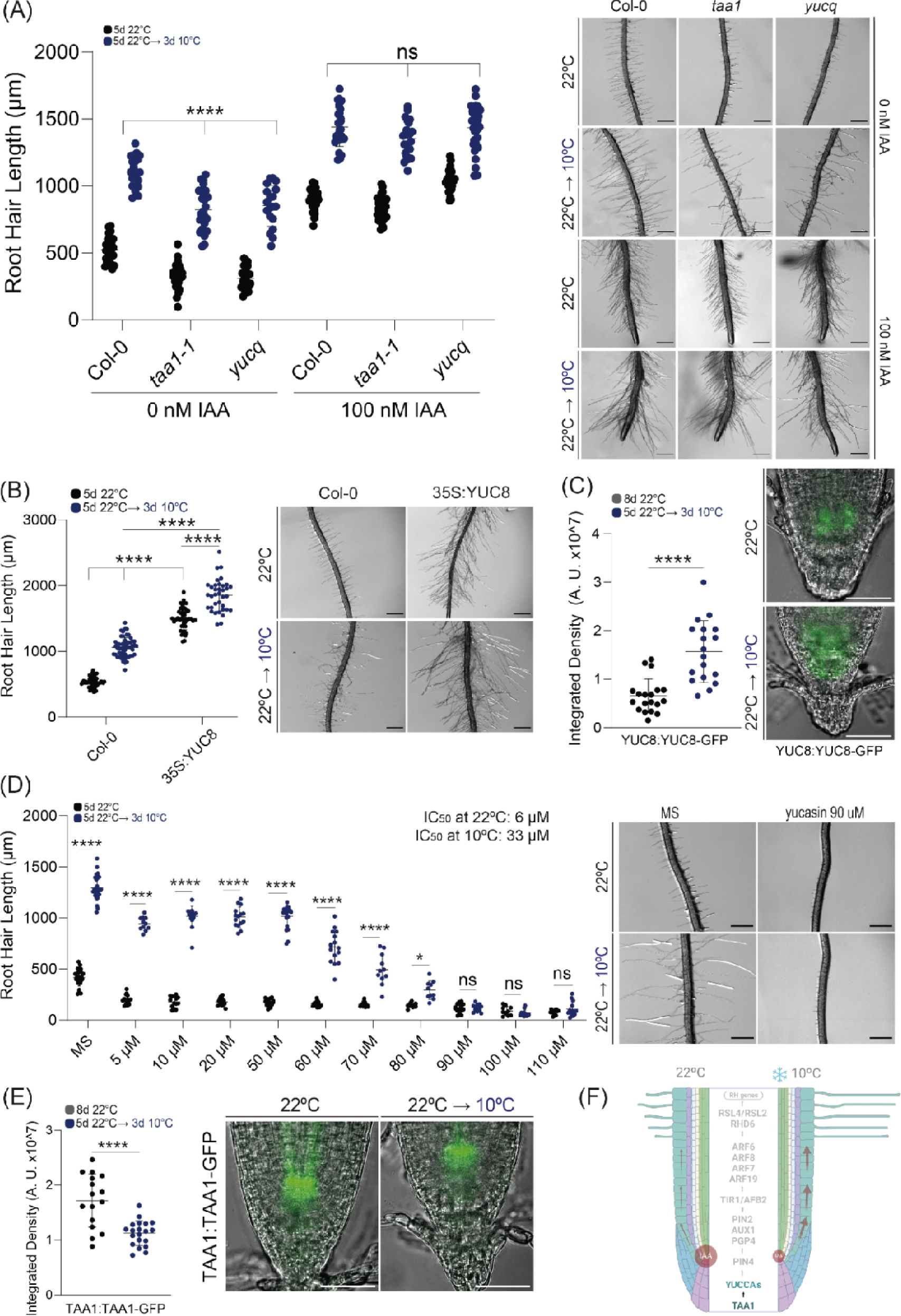
Auxin biosynthesis is required for RH growth at moderate low temperature. (**A**) Scatterplot of RH length of Col-0, *yucq* (lacking *yuc3, yuc5, yuc7, yuc8* and *yuc9*) and *sav3-3 (taa1)* grown at 22°C or at 10°C and treated with 100 nM IAA or without auxin as control. RH growth is enhanced at low temperature in Wt Col-0 and *yucq* and *sav3-3* mutant. Each point is the mean of the length of the 10 longest RHs identified in the maturation zone of a single root. Data are the mean ± SD (n= 10 roots), two-way ANOVA followed by a Tukey–Kramer test; (****) p <0.001, NS= non-significant. Results are representative of three independent experiments. Asterisks indicate significant differences between Col-0 and the corresponding genotype at the same temperature or between the same genotype at different temperatures. Representative images of each genotype are shown on the right. Scale bars= 500 µm. (**B**) Scatterplot of RH length of Col-0 and YUC8 OE grown at 22°C or at 10°C. RH growth is enhanced at moderate low temperature in Wt Col-0 and YUC8 OE. Each point is the mean of the length of the 10 longest RHs identified in the maturation zone of a single root. Data are the mean ± SD (n= 10 roots), two-way ANOVA followed by a Tukey–Kramer test; (****) p<0.001. Results are representative of three independent experiments. Asterisks indicate significant differences between Col-0 and the corresponding genotype at the same temperature or between the same genotype at different temperatures. Representative images of each genotype are shown on the right. Scale bars= 500 µm. (**C**) Confocal images of the root apex of a fluorescent translational reporter line of YUC8-GFP at 22°C and after transfer from ambient to low temperature (22 → 10°C). Fluorescence intensity is expressed in arbitrary units (AU), n= 20 roots. On the right, the GFP-signal is quantified. Results are representative of three independent experiments. Scale bars= 50 μm. (**D**) Scatterplot of RH length of non-treated and treated Col-0 with yucasin (inhibitor of Yucca activity) grown at 22°C or at 10°C. RH growth is enhanced at low temperature in Wt Col-0. Each point is the mean of the length of the 10 longest RHs identified in the maturation zone of a single root. Data are the mean ± SD (n= 10 roots), two-way ANOVA followed by a Tukey–Kramer test; (****) p<0.001, NS= non-significant. Results are representative of three independent experiments. Asterisks indicate significant differences between Col-0 and the corresponding treatment at the same temperature or between the same treatment at different temperatures. Representative images of each treatment are shown on the right. Scale bars= 500 µm. (**E**) Confocal images of root apex of a fluorescent translational reporter line of TAA1-GFP at 22°C and after transfer from ambient to low temperature (22 → 10°C). Fluorescence intensity is expressed in arbitrary units (AU), n= 20 roots. On the right, the GFP-signal is quantified. Results are representative of three independent experiments. Scale bars = 50 μm. (**F**) Summary of the results obtained.

**Figure 3.**
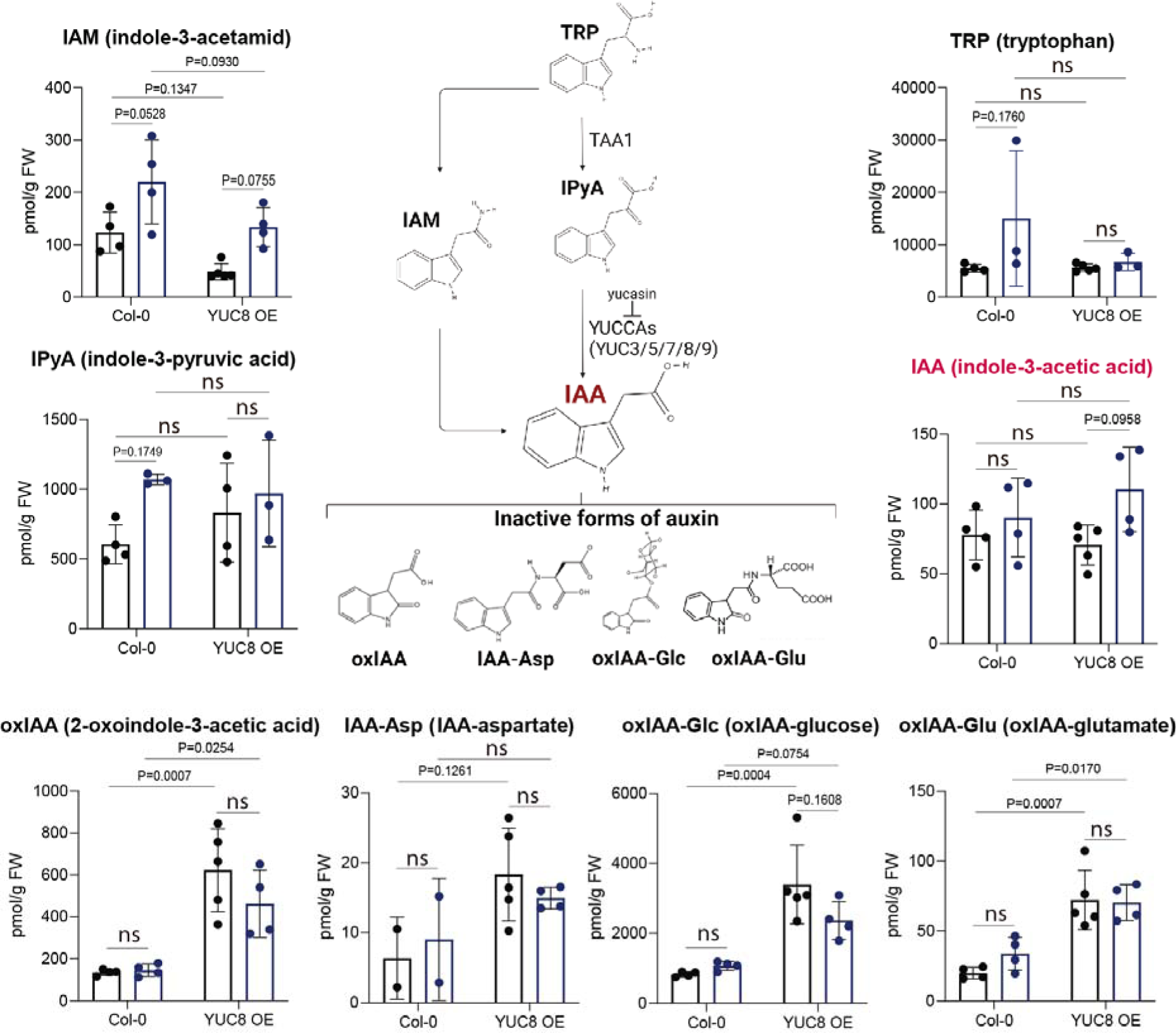
Chemical changes in the Auxin-related pathway linked to moderate low temperature. Auxin precursors (TRP, Tryptophan; IPyA, indole-3-pyruvic acid; IAM, indole-3-acetamid), and IAA (indole-3-acetic acid) and inactive forms (oxIAA, 2-oxindole-3-acetic acid, IAA-Asp, IAA-aspartate; oxIAA-Glc, oxIAA-glucose; oxIAA-Glu, oxIAA-glutamate) were identified and quantified by high-performance liquid chromatography coupled with mass spectrometry (UHPLC-MS/MS) in Wt Col-0 and YUC8 OE line from roots grown at 22°C or at 10°C. Data are the mean ± SD (N= roots), two-way ANOVA. Only P-values below 0.2 are indicated. Results are representative of three independent determinations. See also **Table S2** for the quantitative determinations.

### Auxin transport is mediated by PIN2, PIN4 and AUX1, which affect on RH growth at low-temperature

To investigate how moderate low temperature induced auxin increase is sustained in root apical meristems, we examined the role of polar auxin transport in more detail by assessing the roles of PINs and AUX1, which mediate shoot-ward auxin transport from the root tip to the epidermis in the elongation and differentiation zones of roots (Swarup *et al*., 2012; Wiśniewska *et al*., 2006). As expected, RH elongation of *aux1*, *eir-1-1* (*pin2*) and *pgp4-1* mutants under low temperature was significantly lower than the WT (**Figure 4**). In addition, *pin4* also showed a lack of low temperature RH response. In agreement with this, both PIN2 and PIN4 were upregulated under low temperature in the RNA-seq gene expression profiling (**Figure 1**). Before it was demonstrated that AUX1 and PIN2 contribute to most of the total RH auxin uptake capacity (Dindas *et al*., 2018). Hence, in the absence of auxin uptake carrier activity in *aux1*, the IAA-dependent RH elongation response to low Pi conditions is severely perturbed. By adding 100 nM of auxin (IAA, indole acetic acid) exogenously, *aux1* and *pgp4-1* mutants were restored to WT Col-0 RH growth while *pin4* and *pin2* did not restore their growth at moderate low temperature (**Figure 4A**). Furthermore, the addition of NOA (1-naphthoxyacetic acid) as an inhibitor that blocks the activities of both *auxin* influx and efflux carriers (Parry et al. 2001), the RH growth is inhibited specifically at moderate low temperature (**Figure 4B**). At the same time, only PIN2:PIN2-GFP, but not PIN4:PIN4-GFP and AUX1:AUX1-GFP protein reporter, increase their signals under low temperature (**Figure 4C-E**). In agreement, PIN2 accumulation in the lateral root cap (LRC) and epidermis is crucial for RH elongation (Swarup *et al*., 2007; Jones *et al*., 2009). AUX1 level decreases under low temperature, which might indicate an unknown posttranslational regulation. Collectively, these results demonstrate that TAA1/YUC8-mediated local auxin biosynthesis in the root apex, PIN4 controls local transport close to the root tip, and AUX1-PGP4/PIN2-driven auxin shootward transport via epidermal tissues is indispensable to stimulate RH elongation in response to low temperature (**Figure 4F**). To test if there are modified levels of auxin for the enhanced RH growth, we used DII-VENUS that reflects the nuclear auxin signaling and distribution (Brunoud *et al*., 2012) and DR5:GFP auxin reporter (Heiser et al.2005) in both the root meristematic and differentiation zones (**Figure 5A-B**). Confocal imaging revealed that auxin response was significantly lower in root epidermal cells of WT under low temperature, visualized by DR5-GFP. In agreement, DII-Venus showed an enhanced signal in the meristematic zone. We demonstrate that low temperature modifies polar auxin transport and nuclear auxin signaling to regulate RH growth (**Figure 5C**).

**Figure 4.**
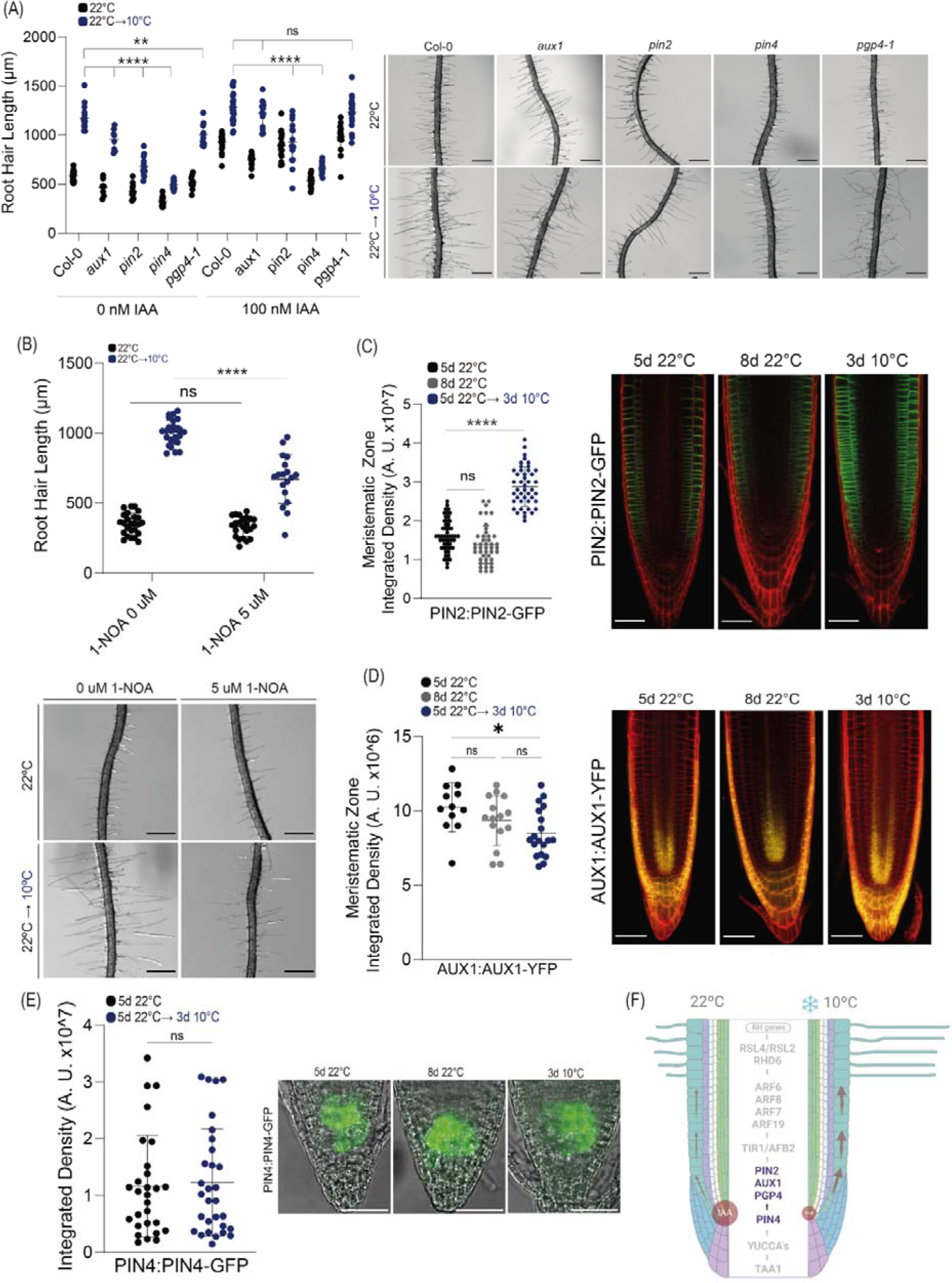
Auxin transport mediated by PIN2/PIN4 and AUX1/PGP4 triggers RH growth at moderate low temperature. (**A**) Scatterplot of RH length of Col-0, *aux1, eir1-1* (*pin2*), *pin4* and *pgp4-1* grown at 22°C or at 10°C. RH growth is enhanced at low temperature in Wt Col-0. Each point is the mean of the length of the 10 longest RHs identified in the maturation zone of a single root. Data are the mean ± SD (n= 20 roots), two-way ANOVA followed by a Tukey–Kramer test; (****) p <0.001, (**) p <0.01. Results are representative of three independent experiments. Representative images of each genotype are shown on the right. Scale bars= 500 µm. (**B**) Scatterplot of RH length of non-treated and treated Col-0 with 1-NOA (auxin influx and efflux inhibitor) grown at 22°C or at 10°C. (**C**) Confocal images of the root apex of a fluorescent translational reporter line of PIN2 (PIN2:PIN2-GFP) at 22°C and after transfer from ambient to low temperature (22→10°C). Fluorescence intensity is expressed in arbitrary units (AU), n=20 roots. On the right, GFP-signal is quantified. Results are representative of three independent experiments. Scale bars = 50 μm. (**D**) Confocal images of the root apex of a fluorescent translational reporter line of AUX1 (AUX1:AUX1-YFP) at 22°C and after transfer from ambient to low temperature (22→10°C). Fluorescence intensity is expressed in arbitrary units (AU), n=20 roots. On the right, YFP-signal is quantified. Results are representative of three independent experiments. Scale bars = 50 μm. (**D**) Confocal images of root apex of a fluorescent translational reporter line of PIN4 (PIN4:PIN4-GFP) at 22°C and after transfer from ambient to low temperature (22→10°C). Fluorescence intensity is expressed in arbitrary units (AU), n=20 roots. On the letf, the GFP-signal is quantified. Results are representative of three independent experiments. Scale bars = 50 μm. (**F**) Summary of the results obtained.

**Figure 5.**
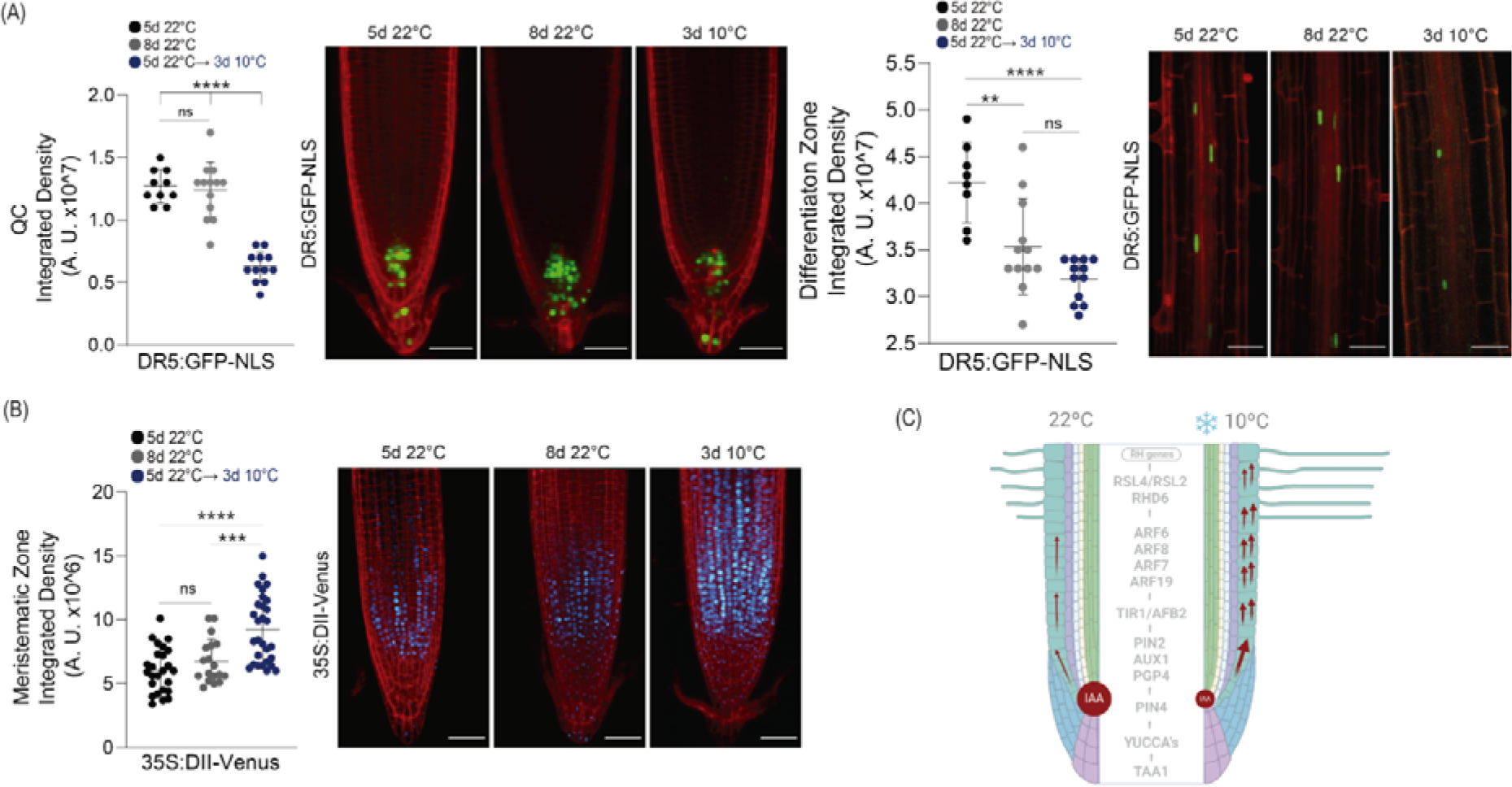
Effect of moderate low temperature on the auxin signaling reporters DR5-GFP and DII-venus. (**A**) Quantitative evaluation of DR5:GFP-NLS reporter fluorescence intensity across the root elongation zones at 22°C and after transfer from ambient to moderate low temperature. Fluorescence intensity is expressed in arbitrary units (A.U.), n=20 roots, 10 root hairs. Results are representative of three independent experiments. (****) p <0.001, (**) p <0.01, (*) p <0.05. On the bottom, confocal images of DR5:GFP-NLS reporter line of at 22°C and after transfer from ambient to low temperature (22°C→10°C). Scale bars=5 μm. (**B**) Quantitative evaluation of 35S:DII-Venus reporter fluorescence intensity across the root differentiation zones at 22°C and after transfer from ambient to low temperature. Fluorescence intensity is expressed in arbitrary units (A.U.), n=20 roots, 10 root hairs. Results are representative of three independent experiments. (****) p <0.001, (***) p <0.005. (**C**) Summary of the results obtained.

### Auxin TIR-AFBs and ARFs mediated signaling effect on RH growth at low temperature

To cause RH swelling and RH growth, the TRANSPORT INHIBITOR1 (TIR1)/AUXIN SIGNALING F-BOX (AFBs) and Aux/IAA co-receptors need first to detect auxin in the trichoblasts. Based on this, we tested if *tir1-1* and the double mutant *tir1 afb2-3* showed any defect in RH growth under low temperature. TIR1 was already detected in the RNA-seq experiment (**Figure 1**). Both *tir1-1* and the double mutant *tir1 afb2-3* showed much shorter RHs than WT at moderate low temperature while the effect was minor at 22°C (**Figure 6A**). The *tir1 afb2,4,5* multiple mutant showed a similar phenotype than *tir1-1* and the double mutant *tir1 afb2-3*. By adding 100 nM of IAA exogenously, both mutants were restored to WT Col-0 RH growth (**Figure S1**), suggesting that multiple AFBs may be acting together with TIR1. Interestingly, TIR1:TIR1-GFP showed no clear change in the expression level under low temperature in the expansion and differentiation zones where RH develops (**Figure 6B**). Upon arrival to the RH zone, auxin triggers a combination of short-and long-term signaling events coordinated by ARF transcription factors (Velasquez *et al*., 2016; Dindas *et al*., 2018). To explore how downstream auxin signaling regulates low temperature-induced RH elongation, we characterized RH phenotypes of a set of several loss-of-function mutants for root-expressed ARF genes (Rademacher *et al*., 2011). Low temperature-induced RH elongation was significantly impaired in the *arf6/arf8*, and *arf7/arf19* single mutants (**Figure 6C**). A further decrease in RH length specifically at low temperature was observed in the *arf6 arf8* double mutant compared to the corresponding single mutants. All suggest that ARF6/ARF8 and ARF7/ARF19 additively regulate the RH elongation under low temperature. In agreement, the ARF7:ARF7-GFP reporter showed enhanced expression at low temperature, and specifically, in epidermals cells, including RHs (**Figure 6D-E**). In agreement with this, all these four ARFs were upregulated under moderate low temperature in the RNA-seq gene expression profiling (**Figure 1**). These results together indicate that at least these four ARFs (ARF6/ARF8 and ARF7/ARF19) are required to trigger RH growth under low temperature (**Figure 6F**).

**Figure 6.**
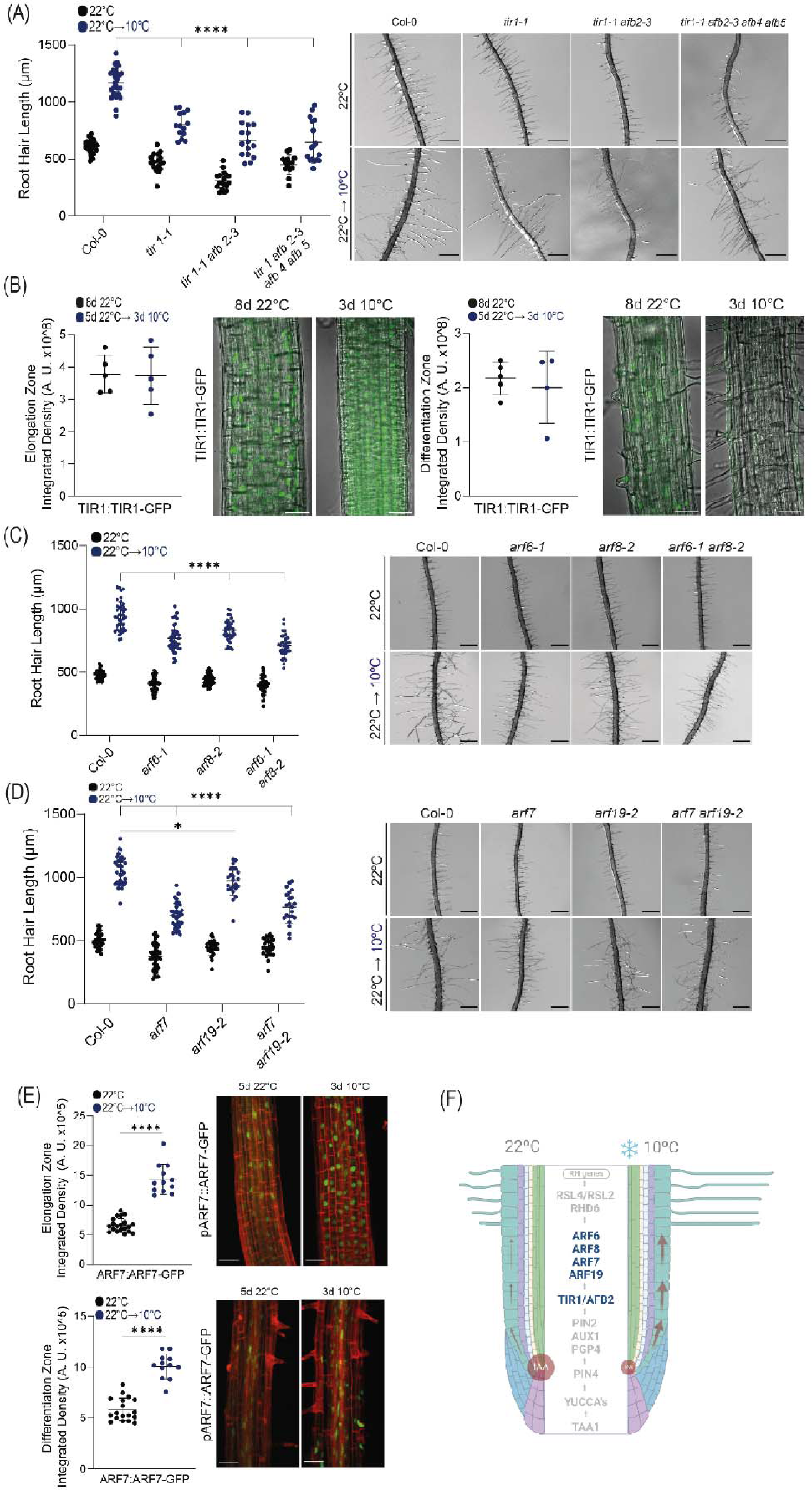
TIR1/AFB2 and multiple ARFs mediate auxin signaling effect on RH growth at moderate low temperature. (**A**) Scatterplot of RH length of Col-0, *tir1-1,* and *tir1-1 afb2,4,5* multiple mutant grown at 22°C or at 10°C. Each point is the mean of the length of the 10 longest RHs identified in the maturation zone of a single root. Data are the mean ± SD (n=20 roots), two-way ANOVA followed by a Tukey–Kramer test; (****) p <0.001, NS= non-significant. Results are representative of three independent experiments. Representative images of each genotype are shown on the right. Scale bars= 300 µm. (**B**) Confocal images of TIR1 translational reporter line of at 22°C and after transfer from ambient to low temperature (22°C→10°C) On the bottom: semi-quantitative evaluation of the GFP fluorescence intensity across the subapical root hair zone at 22°C and after transfer from ambient to low temperature, indicated by a interrupted white line. Fluorescence intensity is expressed in arbitrary units (A.U.), n=20 roots, 10 root hairs. Results are representative of three independent experiments. Scale bars=5 μm. (**C**) Scatterplot of RH length of Col-0, *arf6-1*, *arf8-2* and double mutant *arf6-1 arf8-2* grown at 22°C or at 10°C. RH growth is enhanced at low temperature. Each point is the mean of the length of the 10 longest RHs identified in the maturation zone of a single root. Data are the mean ± SD (n=20 roots), two-way ANOVA followed by a Tukey–Kramer test; (****) p<0.001, NS= non-significant. Results are representative of three independent experiments. Representative images of each genotype are shown on the right. Scale bars= 500 µm. (**D**) Scatterplot of RH length of Col-0, *arf7*, *arf 19-2* and double mutant *arf7 arf19-2* grown at 22°C or at 10°C. RH growth is enhanced at low temperature. Each point is the mean of the length of the 10 longest RHs identified in the maturation zone of a single root. Data are the mean ± SD (n=10 roots), two-way ANOVA followed by a Tukey–Kramer test; (****) p<0.001, (*) p<0.05, NS= non-significant. Results are representative of three independent experiments. Representative images of each genotype are shown on the right. Scale bars= 500 µm. (**E**) Confocal images of ARF7:ARF-GFP translational reporter line of at 22°C and after transfer from ambient to low temperature (22°C→10°C). Quantitative evaluation of the GFP fluorescence intensity across the root differentiation zones at 22°C and after transfer from ambient to low temperature. Fluorescence intensity is expressed in arbitrary units (A.U.), n=20 roots, 10 root hairs. Results are representative of three independent experiments. (****) p<0.001. Scale bars=5 μm. (**F**) Summary of the results obtained.

## Discussion

Plant root In highly heterogeneous soil environments require developmental plasticity to optimize resource acquisition. RH elongation is an essential adaptation for acquiring immobile nutrients such as phosphate and nitrates (Yi *et al*., 2010; Bhosale *et al*., 2018; Jia *et al*., 2023). Here, we report that under low temperature conditions, Arabidopsis roots use the hormone auxin to enhance RH growth and trigger uptake of essential nutrients. Our study demonstrates that promoting RH elongation under moderate low temperature requires the concerted activity of auxin synthesis, transport, and transcription-related components, allowing us to construct a mechanistic framework for this essential root adaptive response pathway. Moderate low temperature, as previously indicated, causes more significant RH formation by directly affecting the availability of nitrate (Pacheco *et al*., 2023), and its influence on phosphate and in other macronutrient dynamics in the media/soil has not yet been determined. It is well known that these two crucial macronutrients are first noticed at the root tip and that a lack of them causes the development of RH (**Figure 7**). Auxin production, transport, and signaling, which depend on both nutrients, convey the signal from the root tip to the differentiation zone, where RH develops. Moderate low temperature possibly triggered low levels of phosphate and nitrate are initially seen at the root tip due to increased expression of TAA1 and YUCCA genes, which causes high auxin levels in the root tip (Bhosale *et al*., 2018; Jones *et al*., 2009; Jia *et al*., 2023; this study). The additional auxin produced at the root tip must either be inactivated as oxIAA, oxIAA-Glu and oxIAA-Glc or reach the RHs for them to grow by the basipetal movement of auxin (shoot-wards). Low temperature triggered at the plasma membrane, higher auxin transport regulated first by PIN4 in the root tip, and then by higher levels of PIN2 at the root epidermis layers (Gangully *et al*., 2010; Giri *et al*., 2018). However, the mechanisms of their inductions may not be only based on transcriptional activation (**Figure 7**). It’s still possible that an enhanced symplastic route of Auxin flow through plasmodesmata (Mellor et al., 2020) might be also involved in RH growth but it was not tested here. To cause RH swelling and development, TIR1/AFB2 (and possible other AFBs) were involved in the auxin sensing in the moderate low temperature conditions in the T cells before RH emerged (**Figure 7**). Downstream, ARF7/ARF19 and ARF6/ARF8 are the ARFs that are involved in RH growth response under low temperature. ARF7/ARF19 might encourage PHR1 expression and are PHR1’s primary targets (Huang et al. 2018) while the two most important ARFs for regulating the RH growth response in the presence of low nitrate are ARF6 and ARF8, according to a recent study (Jia *et al*., 2023). These two groups of ARFs positively control the expression of numerous transcription factors, including ROOT HAIR DEFECTIVE-SIX LIKE 2 (RSL2), RSL4 (Yi *et al*., 2010; Mangano *et al*., 2017; Marzol et al. 2017; Bhosale *et al*., 2018), and *Lotus japonicus* ROOT HAIRLESS-LIKE 3 (LRL3) (Karas et al. 2009), as well as other crucial components of polar development (Mangano et al. 2017; Zhu et al. 2020). RHD6, along with its partially redundant homolog RSL1, is a fundamental helix-loop-helix transcription factor (Menand et al. 2007; Takeda et al. 2008) that works as a master regulator to enhance the expression of its downstream transcription factors RSL2, RSL4, and LRL3. RSL4/RSL2 are the TFs that react to low phosphate levels (Bhosale et al. 2018), while RHD6/LRL3 are the TFs that respond to low nitrate levels (Jia *et al*., 2023). Low phosphate and low nitrate sensing diverge at the downstream transcriptional targets of RHD6, where either RSLs or LRLs take over (Jia et al. 2023; Lopez et al. 2023). On the contrary, under excess of nutrients, the trihelix transcription factor GT2-LIKE1 (GTL1) and its closest homolog, DF1, are able to stop RH growth by directly repressing RSL4 and RSL4 target genes (Shibata et al., 2018; Shibata et al., 2022). In addition, GTL1 is able to bind to RHD6 and prevent the activation of RSL4 (Shibata et al., 2022). Overall, auxin and the downstream activation-repression establish an equilibrium of the intensity of RH growth is being triggered depending on the nutritional contexts that is affected by changes in the temperature of the media-soil (Datta et al., 2015; Bhosale et al.2018; Giri et al. 2018; Jia et al. 2023). Moderate low temperature stimulus on root and RHs comprises complex nutritional signals coming from the media-soil.

**Figure 7.**
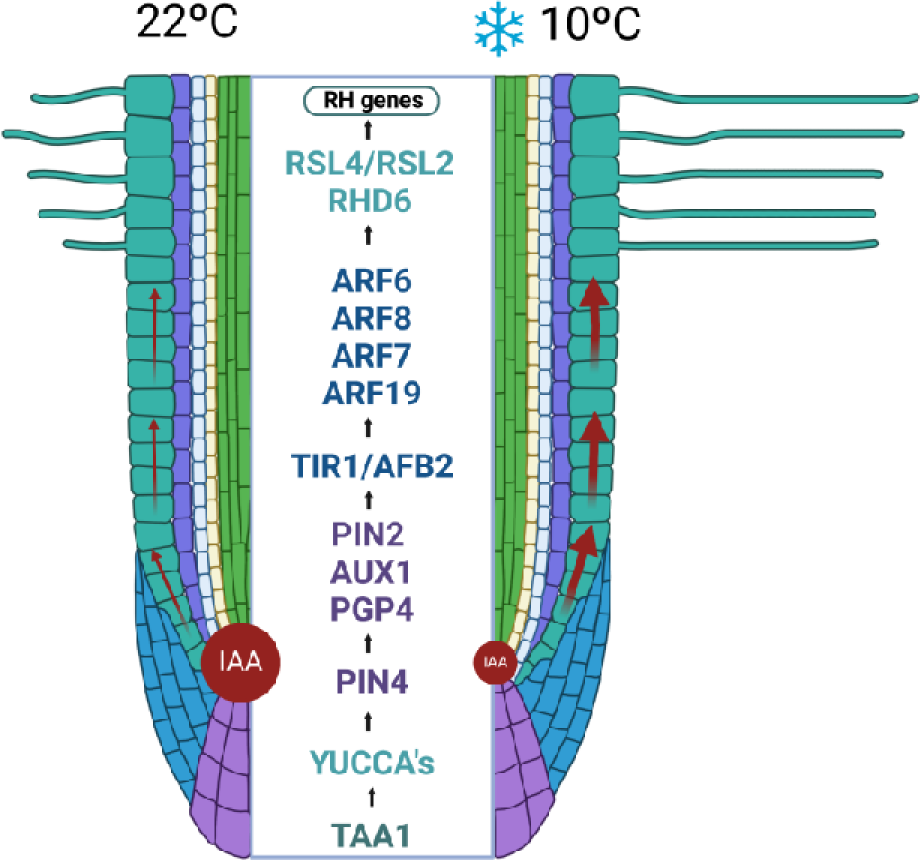
Proposed model on how moderate low temperature affects the auxin pathway in the root that impacts on root hair (RH) growth. This model proposes a regulatory mechanism starting in the root tip to control the foraging response of RH under moderate low temperature. The levels of TAA1 and YUC8 transcripts and YUC8 protein rise in the root apical meristem, resulting in higher amounts of auxin in the root apex. PIN4 then transports auxin out of the tip of the meristem. Then, auxin accumulating in the local area is carried towards the shoot via AUX1 and PIN2/PGP4 proteins in the epidermal cells, specifically to the zone where elongation and differentiation occur. PIN2 expression is enhanced. Auxin is then perceived by TIR1 and AFB2 (by other AFBs) in trichoblast cells and then a transcriptional activation of RH-specific genes by four ARFs (ARF6/ARF8 and ARF7/ARF19) and a downstream RHD6-RSL4/RSL2 cascade (Moison et al. 2021), leading to the intense stimulation of RH elongation.

In summary, our work has revealed that moderate low temperatures directly influence auxin synthesis, transport, and signaling, affecting nutrient mobility, both of which impact root hair (RH) growth. Other hormones, such as ethylene, cytokinin, and strigolactones, may also play a role in the interaction between auxin and nutrient dynamics (recently reviewed in Lopez et al., 2023). Enhancing RH density and length could be a cost-effective strategy to improve nutrient acquisition, especially for soil nutrients with low mobility, ultimately boosting plant fitness. Our findings, in conjunction with other pivotal studies on RH growth (Yi et al., 2010; Bhosale et al., 2018; Giri et al., 2018; Jia et al., 2023), lay the foundation for developing a molecular framework aimed at optimizing root nutrient foraging. This could lead to identifying novel breeding targets for more efficient nutrient-uptake configurations in crops.

## Experimental Procedures

### Plant Material and Growth Conditions

Arabidopsis (*Arabidopsis thaliana*) Columbia-0 (Col-0) ecotype was used as a wild-type genotype in all experiments. All mutants and transgenic lines tested are in this ecotype. Seedlings were germinated on agar plates in a Percival incubator at 22°C in a plant growth chamber in continuous light at 120 μmol.seg^−1^.m^−2^ intensity for 5 days at 22°C + 3 days at 10°C for the low temperature response. Plants were transferred to soil for growth under the same conditions as previously described at 22°C. For identification of T-DNA knockout lines, genomic DNA was extracted from rosette leaves. Confirmation by PCR of a single and multiple T-DNA insertions in the target genes were performed using an insertion-specific LBb1 or LBb1.3 (for SALK lines) or Lb3 (for SAIL lines) primer in addition to one gene-specific primer. To ensure gene disruptions, PCR was also run using two gene-specific primers, expecting bands corresponding to fragments larger than in Wt. In this way, we isolated homozygous lines (for all the genes in this study). For the imaging of fluorescence intensity distribution in root tips and RH, seedlings were grown for 5 days at 22°C plus 3 days at 10°C. All mutants and transgenic lines are listed in **Table S4**.

### Pharmacological Treatments

For all experiments, plants were grown first on solid 0.5X MS medium at 22°C for 3 days in continuous light. According to the specific treatment plants were transferred to plates containing regular solid 0.5X MS supplemented with 100nM IAA (auxin treatment), 5 μM 1-NOA (1-Naphthoxyacetic acid, M.W. 186.21) or 5-110 μM Yucasin (5-(4-chlorophenyl)-4H-1,2,4-triazole-3-thiol, M.W. 211.67) and grown at 22°C for 3 days plus 3 days at 10°C in continuous light. RH phenotype was quantified after each 3 day span. Yucasin and 1-NOA were dissolved in DMSO.

### Root hair phenotype

Seeds were surface sterilized and stratified in darkness for 3 days at 4°C. Then grown *in vitro* in a MS 0.5X medium (pH 5.7; 0.8% w/v agar) in a plant growth chamber in continuous light (120 μmol.sec^−1^.m^−2^) at 22°C and/or 10°C. The quantitative analyses of RH phenotypes of Col-0 and transgenic lines were made the last day of the growth conditions described in the two previous sections. In the case of low temperature treatment, the measurements were done after 5 days at 22°C and after 3 days at 10°C. For that purpose, 10 fully elongated RH from the maturation zone were measured per root under the same conditions from each treatment and control. 20 roots were measured as indicated in each case. Images were captured using an Olympus SZX7 Zoom Stereo Microscope (Olympus, Tokyo, Japan) equipped with a Q-Colors digital camera and Q Capture Pro 7 software (Olympus, Japan). Results were expressed as the mean ± SD using the GraphPad Prism 8.0.1 (GraphPad Software, Boston, MA, USA) statistical analysis software. Results are representative of three independent experiments, each involving 30 roots.

### Confocal Microscopy

For measurements of fluorescence intensity distributions after cold stress (22°C→10°C) in root tips and root hairs of DR5:GFP-NLS, PIN2:PIN2-GFP, PIN4:PIN4-GFP, AUX1:AUX1-YPF, ARF7:ARF7-GFP, TIR1:TIR1-GFP lines confocal laser scanning microscopy Zeiss LSM 710 (Carl Zeiss, Germany) was used. For image acquisition, 20x/1.0 NA Plan-Apochromat objective for root tips and 40x/1.4 Oil DIC Plan-Apochromat objective for root hairs were used. The GFP signal was excited with a 488 nm argon laser at 4% laser power intensity and emission band of 493-549 nm. Propidium Iodide signal was excited with a 488 nm argon laser at 4% laser power intensity and emission band 519-583 nm. Fluorescence AU was expressed as the mean ± SD using the GraphPad Prism 8.0.1 (USA) statistical analysis software. Results are representative of three independent experiments, each involving 10-20 roots as indicated in each case.

### RNA-seq analyses

For the RNA-seq analysis, seedlings were grown on ½ strength MS agar plates, in a plant growth chamber at 22°C in continuous light (120 μmol s^−1^ m^−2^) for 14 days at 22°C as a pretreatment and then at 10°C (moderate-low temperature treatment) for 0hs, 2hs and 6hs. We analyzed a dataset with 6 factor groups (tree time points and two genotypes: Col-0 and *rsl2rsl4*, each with three biological replicates) giving 18 samples in total. Total RNA was extracted from 20–30 mg of frozen root tissue. Frozen root samples were ground in liquid nitrogen and total RNAs were extracted using E.Z.N.A Total RNA Kit I (Omega Bio-tek, Georgia, USA). RNA quantity and purity were evaluated with a Qubit®2.0 fluorometer (InvitrogenTM, Carlsbad, CA, USA) using a QubitTM RNA BR assay kit. RNA integrity and concentration were assessed by capillary electrophoresis using an automated CE Fragment AnalyzerTM system (Agilent Technologies, Santa Clara, CA, USA) with the RNA kit DNF-471-0500 (15nt). Total RNA-seq libraries were prepared according to the TruSeq Stranded Total RNA Kit (Illumina, San Diego, CA, USA) following the manufacturer’s instructions. Finally, the constructed libraries were sequenced using Macrogen sequencing services (Seoul, Korea) in paired end mode on a HiSeq4000 sequencer. For total RNA differential expression analysis,a quality check was performed with FASTQC software (Andrews, 2010). Then, the adapter sequences were removed, reads with a quality score less than 30 and length less than 60 nucleotides were eliminated using Flexbar (Dodt et al., 2012). Resulting filtered reads were aligned against *Arabidopsis thaliana* Araport 11 genome with the STAR aligner software. A total of 18 RNA libraries were sequenced, obtaining an average of 84,678,135 reads for each one, with a minimum and maximum value of 72,658,864 and 100,764,020 reads, respectively. After filtering them by quality and removing adapters, an average of 98.8% of the reads remained and after aligning them against the *Arabidopsis thaliana* reference genome, between 88.1% (69,820,848) and 96.5% (93,072,528) of total reads were correctly aligned (**Table S2**). For each library, the feature Counts software from the Rsubread package (Liao et al., 2019) was applied to assign expression values to each uniquely aligned fragment. Differential gene expression analysis was performed using the Bioconductor R edgeR package (Robinson et al., 2010). Differentially expressed genes (DEG) were selected with an FDR<0.05 and a FC>|0.2|. To search for genetic functions and pathways overrepresented in the DEG lists, genetic enrichment analysis was performed using the Genetic Ontology (GO) database with the R package ClusterProfiler v4.0.5 (Yu et al., 2012), using the compareCluster function. The parameters used for this analysis were: lists of differentially expressed genes for each comparison in ENTREZID, enrichGO sub-function, the universe from the total of differentially expressed genes that present annotation as genetic background, Benjamini-Hochberg statistical test and a filter of FDR less than 0.05. Subsequently, the semantics filter of GO terms was performed using the simplify function of the same package using a p-value and q-value cutoff less than 0.05.

### Quantification of IAA metabolites

The determination of endogenous auxin metabolites was conducted following the protocol described by Pěnčík et al. (2018). Briefly, 3-6 samples of approximately 10 mg (fresh weight) for each genotype or condition (roots growth at 22°C for two weeks or 11 days at at 22°C and then transferred for 3 days at 10°C) were freeze-dried and then extracted using 1 mL of cold 50 mmol/L phosphate buffer (pH 7.0) containing 0.1% sodium diethyldithiocarbamate and mixture of stable isotope-labeled internal standards. A 200 µl portion of the extract was acidified to pH 2.7 with HCl and subjected to in-tip micro solid phase extraction (in-tip µSPE). Another 200 µl portion was derivatized with cysteamine, acidified to pH 2.7 with HCl, and purified using in-tip µSPE to determine IPyA. Following elution, samples were evaporated under reduced pressure, reconstituted in 10% aqueous methanol, and analyzed using an HPLC system 1260 Infinity II (Agilent Technologies, USA) equipped with a Kinetex C18 column (50 mmx2.1 mm, 1.7 µm; Phenomenex) and coupled to 6495 Triple Quad detector (Agilent Technologies, USA). See **Table S3** for the quantification details.

## Acknowledgements

We would like to thank all the groups that provide us with the seeds mentioned in **Table S4,** including Mark Estelle, Jiril Friml, Marisa Otregui, Ramiro Paris. We thank NASC (Ohio State University) for providing T-DNA lines seed lines. J.M.E. is an investigator of the National Research Council (CONICET) from Argentina. This work was supported by grants from ANPCyT (PICT2019-0015 and PICT2021-0514), by ANID – Programa Iniciativa Científica Milenio ICN17_022, NCN2021_010, ICN2021_044 (C.M.) and Fondo Nacional de Desarrollo Científico y Tecnológico [1200010] to J.M.E.

## Author Contribution

V.B.G. performed most of the experiments, analyzed the data and helped with the manuscrupt writing process. M.A.I., J.M.P., L.L., M.C. helped in some experiments and in the writing process. G.N.L and C.M. carried out the RNA-seq data analysis. A.P. and O.N. carried out the chemical determination of auxins; Z.J., R.F.H.G and N.vH provided several lines used in this study and helped on the writing process. J.M.E. designed the research, analyzed the data, supervised the project, and wrote the paper. All authors commented on the results and the manuscript. This manuscript has not been published and is not being considered for publication elsewhere. All the authors have read the manuscript and have approved this submission.

---

**Table S4.** Mutants and transgenic lines used and generated in this study.

| Mutants and transgenic lines |  |  |  |  |
| --- | --- | --- | --- | --- |
| Locus | Gene name | Gene expression effect | Line name | Reference |
| Auxin Biosynthesis |  |  |  |  |
| AT1G70560 | TAA1 | mutant | *taa1-1* | SALK_022743C ABRC |
|  |  | reporter | TAA1:TAA1-GFP | Yang et al. 2014 |
| AT1G04610<br>AT5G43890<br>AT2G33230<br>AT4G28720<br>AT1G04180 | *yuc3*<br>*yuc5*<br>*yuc7*<br>*yuc8*<br>*yuc9* | mutant | *yuc quintuple* | Li et al. 2012 |
| AT4G28720 | YUC8 | O.E. | 35S:YUC8 |  |
|  |  | reporter | YUC8:YUC8-GFP | Jia et al. 2023 |
|  |  | reporter | DR5:GFP-NLS | Heiser et al.2005 |
|  |  | reporter | 35S:DII-Venus | Brunoud *et al.*, 2012 |
| Auxin Transport |  |  |  |  |
| AT2G38120 | AUX1 | mutant | *aux1-7* | Pickett et al. 1990 |
|  |  | reporter | AUX1:AUX1-YFP | Jones *et al.*, 2009 |
| AT5G57090 | PIN2 | mutant | *pin2-1* |  |
|  |  | reporter | PIN2:PIN2-GFP | Jones *et al.*, 2009 |
| AT2G01420 | PIN4 | mutant | *pin4-1* |  |
|  |  | reporter | PIN4:PIN4-GFP |  |
| AT2G47000 | PGP4 | mutant | *pgp4-1* |  |
| Auxin signaling |  |  |  |  |
| AT3G62980 | TIR1 | mutant | *tir1-1* |  |
|  |  | reporter | TIR1:TIR1-GFP |  |
| AT3G62980<br>AT3G26810 | TIR1 AFB2 | mutant | *tir1-1 afb2-3* |  |
| AT3G62980<br>AT3G26810<br>AT4G24390<br>AT5G49980 | TIR1<br>AFB2<br>AFB4<br>AFB5 | mutant | *tir1-1*<br>*arfb2-3 afb4-8 afb5-5* |  |
| AT1G30330 | ARF6 | mutant | *arf6-1* | Jia et al. 20123 |
| AT5G37020 | ARF8 | mutant | *arf8-2* | Jia et al. 20123 |
| AT5G37020<br>AT1G30330 | ARF6 ARF8 | mutant | *arf6-1 arf8-2* | Jia et al. 20123 |
| AT5G20730 | ARF7 | mutant | *arf7-1* | CS24607 ABRC |
|  |  | reporter | ARF7:ARF7-GFP |  |
| AT1G19220 | ARF19 | mutant | *arf19-2* |  |
| AT5G20730<br>AT1G19220 | ARF7<br>ARF19 | mutant | *arf7 arf19-2* |  |

